# The impact of multigenerational high-fat diet feeding on the gut microbiome and host metabolism

**DOI:** 10.1101/2022.02.01.478488

**Authors:** Marsha C. Wibowo, Zhen Yang, Theodore A. Chavkin, Brian Nguyen, Loc-Duyen Pham, Tian Lian Huang, Matthew D. Lynes, Yu-Hua Tseng, Aleksandar D. Kostic

**Affiliations:** Joslin Diabetes Center, Section on Pathophysiology and Molecular Pharmacology, Boston, MA, USA; Harvard Medical School, Department of Microbiology, Boston, MA, USA; University of Waterloo, Department of Combinatorics and Optimization, Waterloo, ON, Canada; Massachusetts College of Pharmacy and Health Sciences, Boston, MA, USA; Joslin Diabetes Center, Section on Integrative Physiology and Metabolism, Boston, MA, USA; Maine Medical Center Research Institute, Scarborough, ME, USA; Harvard University, Harvard Stem Cell Institute, Cambridge, MA, USA

## Abstract

Although human genetics partially explain the heritability of obesity and type 2 diabetes (T2D), the human gut microbiome also plays a significant role. While changes in human genetics at the population level occur only after many generations, the gut microbiome evolves over a shorter time span. The gut microbiome is also vertically transmitted across generations, therefore, changes in one generation can be passed on to subsequent generations. However, it is unknown whether the gut microbiome influences natural selection of its host under obesogenic pressure. Here, we show that C57BL/6 mice fed a high-fat diet (60% fat, HFD) over four generations develop resistance to obesity and the metabolic syndrome (MetS). Unexpectedly, the mice were increasingly leaner as well as more glucose tolerant and insulin sensitive across generations. This phenomenon was attributed to the most obese mice not yielding progenies, whereas the leanest mice successfully reproduced, and their offspring were also resistant to obesity. In other words, a population bottleneck was observed. Because all the mice were nearly genetically identical inbred C57BL/6J mice, the large variation in body weight gain in response to HFD feeding was likely independent of genetics. We explored whether microbial factors enriched in obesity-resistant mice promote healthier host metabolic phenotypes under HFD feeding, thereby contributing to the heterogeneity in body weight gain and providing an adaptive advantage to the host. Pearson correlation analysis revealed that body weight gain was positively correlated with *Lactococcus lactis*, as well as negatively correlated with *Lactobacillus johnsonii* and pathways for coenzyme A biosynthesis, amino acid biosynthesis (lysine, isoleucine, valine), and nucleotide biosynthesis (adenosine, guanosine). Overall, we observed multigenerational adaptation in the gut microbiome correlated with improved metabolism, yet further studies are needed to validate that these adaptations drive metabolic health.

## Introduction

Obesity and T2D have become an epidemic. In less than 60 years, the number of Americans with T2D has increased almost 12-fold (CDC), and this number is predicted to continue to rise. In 2015, 36.5% U.S. adults were considered obese^1^, around 9.3% had diagnosed diabetes, 95% of which was T2D, and 90 million Americans were affected by MetS (www.cdc.gov). MetS and diabetes are major drivers of morbidity and mortality, as well as major risk factors for various diseases including renal failure, cardiovascular diseases, and coronary heart disease^2^.

Although human genetics contribute to the heritability of obesity and T2D^3^, the gut microbiome also plays a significant role. Genome-wide association studies have discovered more than 100 loci associated with T2D, but altogether they only explain ~10% of the variation that predisposes someone to T2D^4^. The human gut microbiome consists of trillions of microorganisms residing in our gastrointestinal tract that have the potential to impact virtually every facet of host physiology. It is well established that the gut microbiome plays a causal role in the development of MetS, obesity, and T2D in murine models of these diseases and in humans. For example, fecal microbiota transplantation (FMT) from human twins discordant for obesity transfers lean and obese phenotypes to recipient germ-free (GF) animals^5^. FMT or ‘precision’ probiotic treatment in humans improves insulin sensitivity^6–8^.

The human gut microbiome as a whole is remarkably stable within an individual over long time-spans (decades)^9^, and it is vertically inherited from mother to child^10,11^. However, components of the gut microbiome adjust rapidly in response to environmental stresses, such as a major change in diet^12^. Therefore, components of the microbiome acquired in response to stress that confer an adaptive advantage to their host may be inherited. In this study, we explored the question of whether the gut microbiome influences natural selection of its host under obesity-promoting conditions, such as high-fat diet feeding.

Our results show that C57BL/6 mice fed a high-fat diet (60% fat) were increasingly leaner and more glucose and insulin sensitive across four generations. This phenomenon was due to a population bottleneck in which the most obese mice failed to produce progenies, whereas the leanest mice successfully reproduced. Since all of the mice were nearly genetically identical inbred C57BL/6J mice, the large variation in body weight gain in response to HFD feeding was likely influenced by a non-genetic factor, possibly the gut microbiome. We identified microbial factors that were negatively correlated with weight gain, one of which was *Lactobacillus johnsonii*. We supplemented C57BL/6 mice fed with a HFD with *L. johnsonii* and observed no significant difference in body weight gain compared to controls. Other microbial factors might play a role in influencing the differential survival and reproduction of the mice under a HFD feeding condition.

## Results

### Study design

Previous studies have shown that the metabolic syndrome compounds across generations, in which continuous intake of HFD results in more severe body weight increase across generations (F2>F1>F0)^13–15^, increased lipid accumulation in the liver and white adipose tissue^13,14,16^, increased accumulation of ER stress and lipogenesis in the liver of male mice^15^, and decreased browning of adipose tissue in female mice^13^. Separately, a landmark study by Sonnenburg *et al*. showed that a diet low in MACs (microbiota-accessible carbohydrates) led to progressive loss of gut microbiota diversity across generations^17^. This loss in gut microbiota diversity is irreversible by dietary changes alone without reseeding of ancestral microbiota. Based on these prior studies, we postulated that HFD feeding would result in loss of microbial taxa and worsening of MetS that compound across generations.

To test this, we propagated C57BL/6J mice on a HFD (60% fat, 20% carbohydrate, 20% protein) vs. a chow diet (10% fat, 70% carbohydrate, 20% protein) for four generations **(Fig. 2-1)**. Mice were fed a chow diet for two weeks after weaning and were subsequently divided into experimental (N=10 males) and control (N=8 males) groups. Experimental mice were fed HFD for 10 weeks and reversed to chow for 6 weeks, after which the mice were sacrificed. This reversal to chow was to characterize whether MetS phenotypes and loss of gut microbial diversity could be reversed by dietary intervention. All four generations of mice followed the same dietary treatments. The control group was kept on a chow diet throughout the experiment for four generations. Female mice were not used to avoid stressing mothers and the potential of confounding factors introduced by fluctuation in metabolic phenotypes during pregnancy and nursing. We collected both host and microbiome data longitudinally to understand dynamic alterations in the microbiome and the host.

**Fig. 2-1.**
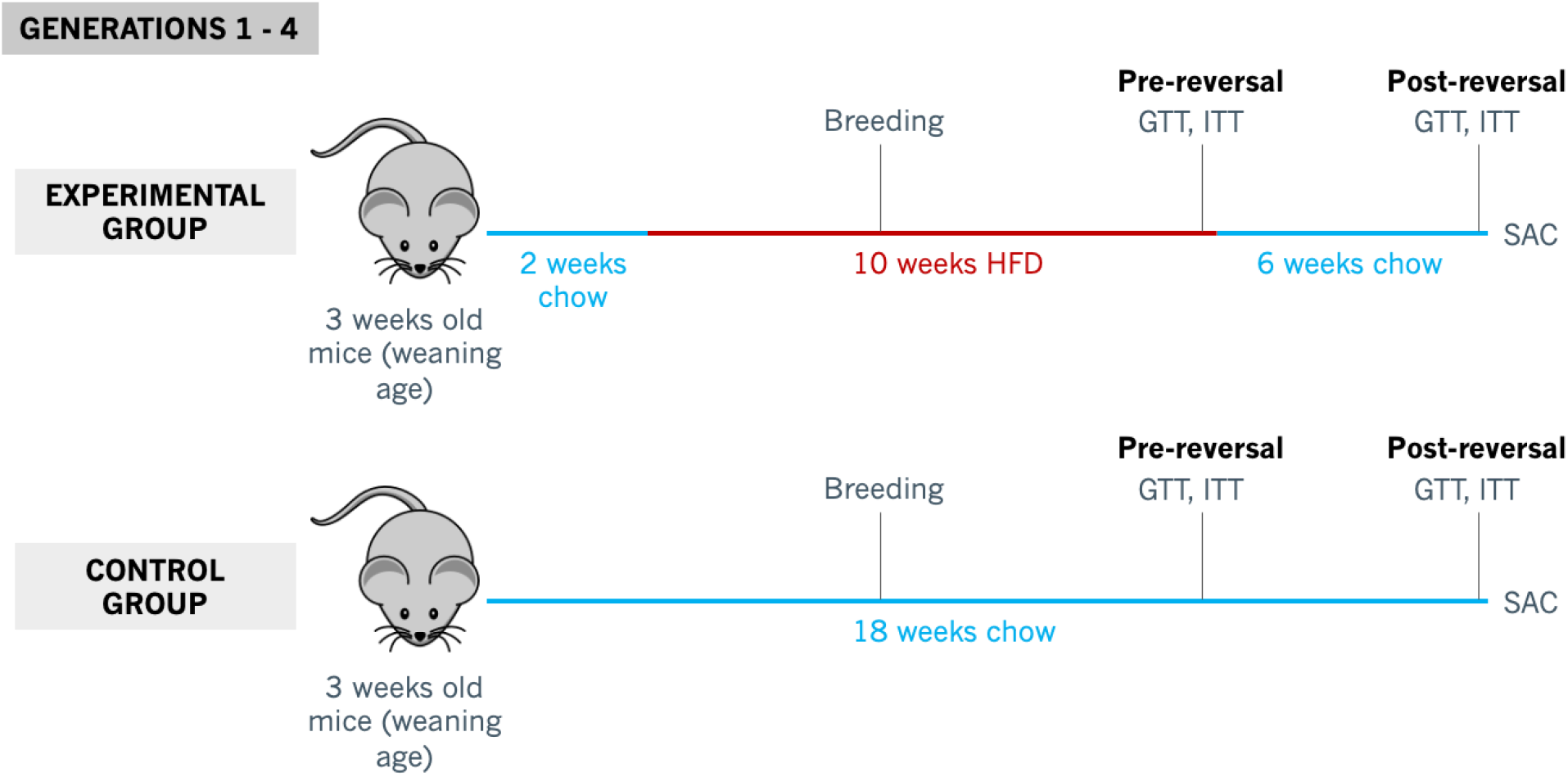
Outline of mouse study design.

### Improved host metabolic phenotypes upon four generations of HFD feeding

To characterize metabolic phenotypes of the host across generations, we measured weekly body weight, food intake, random fed blood glucose levels, and insulin levels. At the end of 10 weeks of HFD (pre-reversal time point) and at the end of 6 weeks of diet reversal to chow (post-reversal time point), we performed intraperitoneal insulin tolerance test (ip-ITT) to measure insulin resistance and intraperitoneal glucose tolerance test (ip-GTT) to quantify glucose intolerance. At the time of sacrifice, we collected blood, pancreas, brown adipose tissue, and major insulin-sensitive tissues (white adipose tissue, muscle, and liver) to assess metabolic changes in the host.

The results revealed that experimental mice from all four generations developed MetS after 10 weeks of HFD feeding and were able to reverse MetS phenotypes after 6 weeks of reversal onto a chow diet **(Fig. 2-2)**. Across all generations, pre-reversal HFD-fed mice had increased body weight gain, as well as worsened glucose tolerance and insulin tolerance relative to chow-fed mice. Reversal of HFD-fed mice onto a chow diet for 6 weeks improved MetS phenotypes. At the post-reversal time point, experimental mice had comparable body weight, glucose tolerance, and insulin tolerance to control mice.

**Fig. 2-2.**
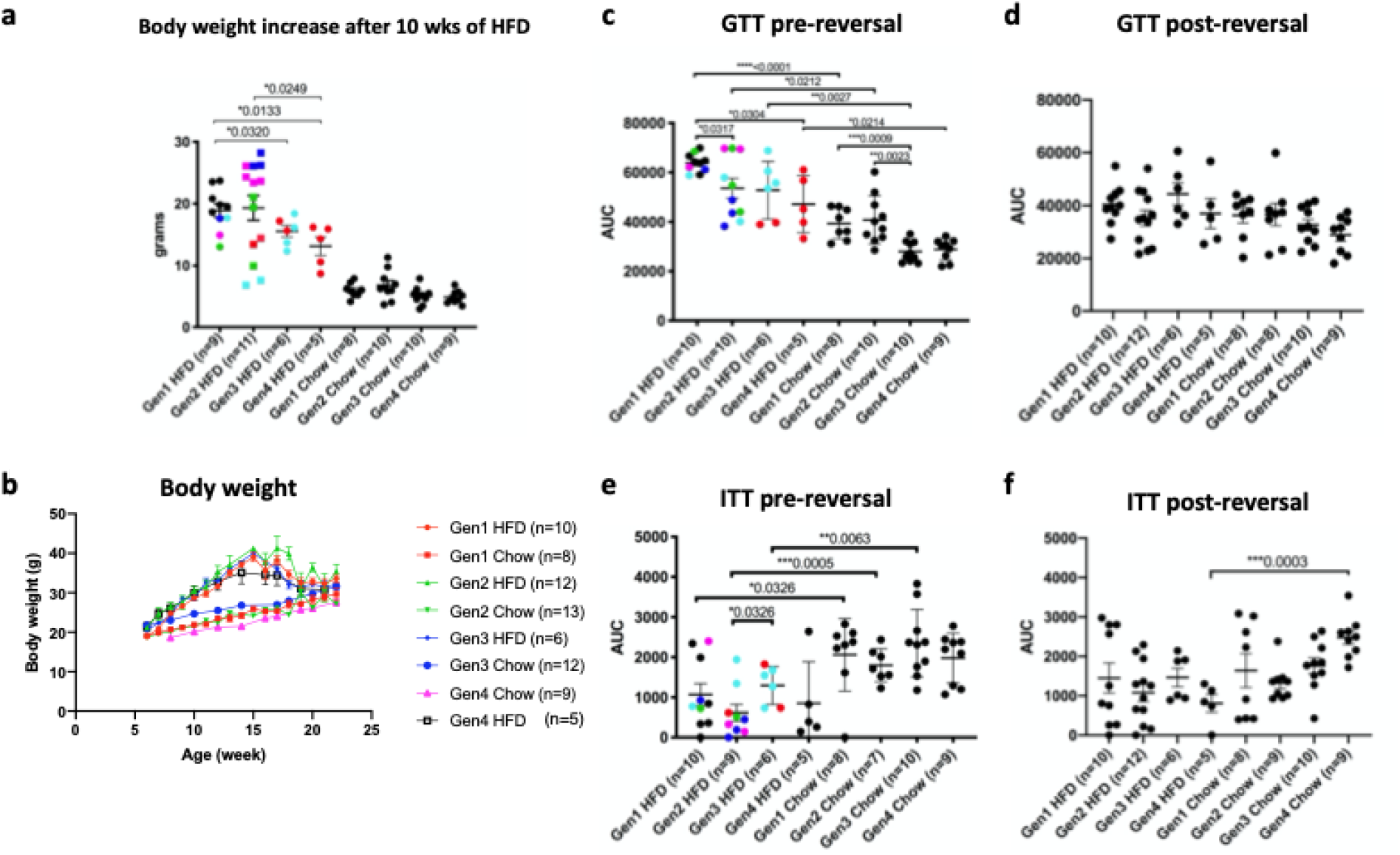
Metabolic syndrome characterization. **a,** Body weight increase after 10 weeks of HFD feeding. **b,** Body weight of the mice up to 22 weeks of age, weekly sampling. **c-d,** Intraperitoneal glucose tolerance test (ip-GTT) on mice after 10 weeks of HFD/pre-reversal (**c**) and after 6 weeks of chow reversal/post-reversal (**d**). **e-f,** Intraperitoneal insulin tolerance test (ip-ITT) on mice pre-reversal (**e**) and post-reversal (**f**). Data are means ± SEM. Significance was computed using two-tailed Welch’s t-test. ns=p-value>0.05; *0.01≤p-value≤0.05; **0.001≤p-value≤0.01; ***p-value<0.001. For Figs. a, c, and e, each point was colored according to each mouse’s paternal lineage.

Contrary to our expectation that 10 weeks of HFD feeding would lead to worsened MetS phenotypes across generations, generation 3 and 4 mice gained less weight relative to generation 1 mice **(Fig. 2-2a)**. Generation 2, 3, and 4 mice were more glucose sensitive compared to generation 1 mice **(Fig. 2-2b)**. Insulin sensitivity did not change significantly relative to generation 1 mice **(Fig. 2-2c)**. This unexpected improvement of MetS phenotypes across generations was likely the result of lower breeding capacity of generation 2 mice that gained significant weight upon HFD feeding (obesity-sensitive mice) relative to the mice that did not gain much weight (obesity-resistant mice) **(Fig. 2-3)**. As a consequence, generation 3 and 4 mice were all offspring of the obesity-resistant parents and were resistant to obesity themselves.

**Fig. 2-3.**
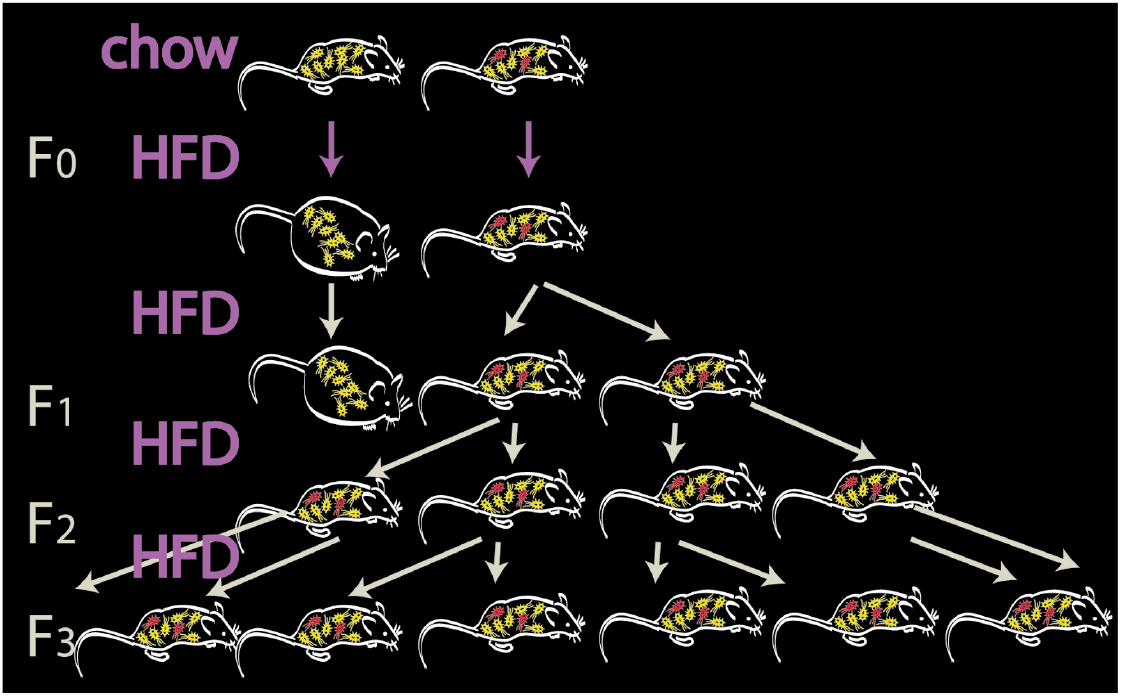
Schematic of hypothesis. A difference in the baseline microbiome drives heterogeneity in glucose control and weight gain in mice. Because the microbiome is inherited and because these phenotypes impact reproductive fitness, natural selection favors mice carrying the beneficial microbiome features (red). However, selection can alternatively be driven by a detrimental microbiome feature in the obesity-sensitive mice.

This result is in contrast to previous studies^13–16^, in which HFD feeding led to worsening of MetS phenotypes across generations. This difference may be due to the fact that in those studies, the fat percentage of the HFD was lower^14,16^ or only one of the parents were fed with a HFD^13,15^. In this study, we fed both parents with a HFD with 60% fat content for 4 generations, which is a stronger selective pressure compared to previous work. We reasoned that under this selective pressure, only mice with increased fitness and better metabolic adaptation to HFD feeding would be able to reproduce **(Fig. 2-3)**. Since C57BL/6 mice are almost free of genetic differences, this variation in susceptibility to obesity is likely due to non-genetic factors. Here, we focused our analysis on whether adaptations in the gut microbiome could explain increased fitness, though other explanations such as epigenetic changes are also possible.

### Microbial species associated with HFD-induced body weight gain

To identify microbial factors that confer host resistance or sensitivity to HFD, we performed shotgun metagenomic sequencing, which allows the identification of unique and shared genes, pathways, and taxonomies between the different groups. We determined species richness in the gut microbiome by running MetaPhlAn2^18^ on the raw reads and counting the number of species per sample. Within the experimental group, there was no significant difference in the average number of species per sample across four generations **(Fig. 2-4a)**. There was also no significant difference in species richness between the experimental group and the control group in each generation.

**Fig. 2-4.**
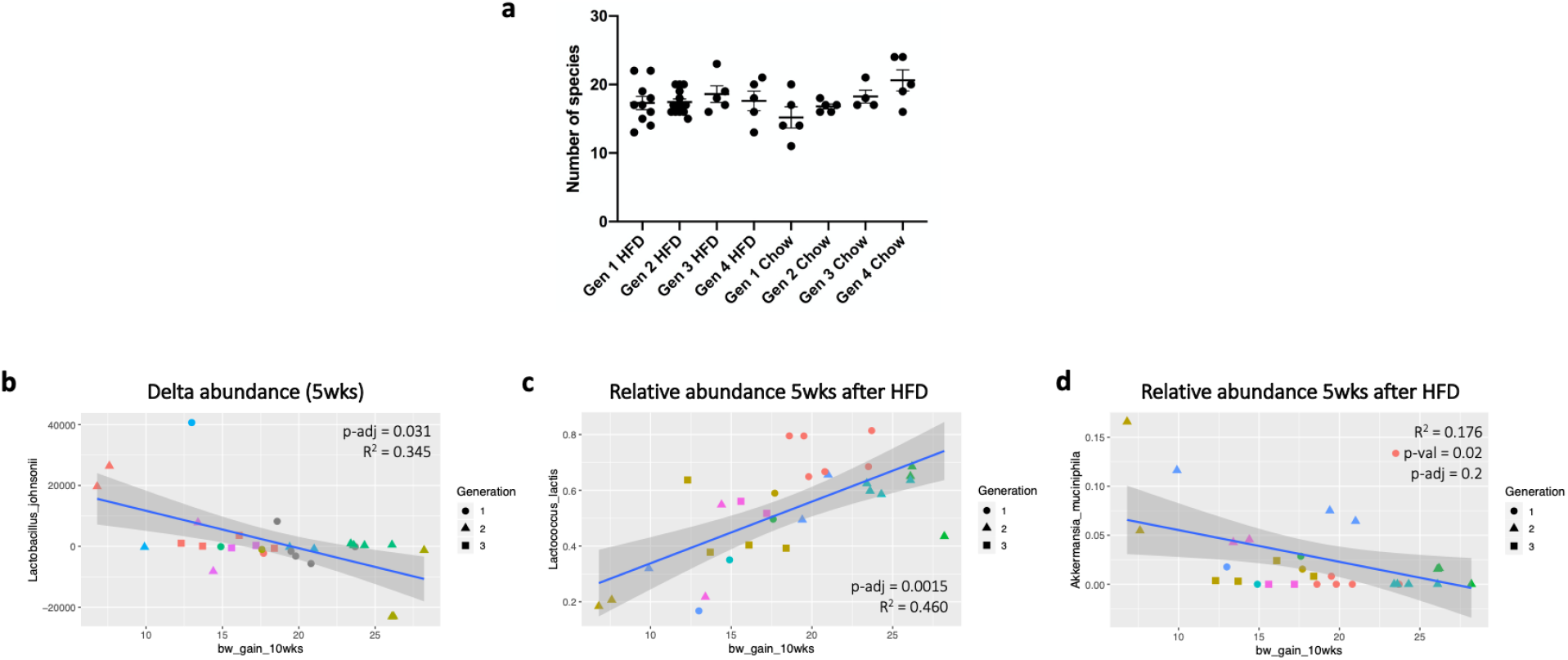
Taxonomic analysis. **a,** Number of species per mouse as identified by MetaPhlAn2. **b-d,** Microbial species correlated with body weight gain. Species was identified using MetaPhlAn2^18^ and Pearson correlation analysis was performed. P-values were corrected with the FDR method.

To identify species associated with body weight gain, we annotated species present in each mouse gut microbiome with MetaPhlAn2^18^ and calculated Pearson correlation coefficients. The results revealed that *Lactococcus lactis* is positively correlated with body weight gain, whereas *Lactobacillus johnsonii* and *Akkermansia muciniphila* are negatively correlated with body weight gain **(Fig. 2-4b)**, although the correlation between weight gain and *A. muciniphila* is not statistically significant after FDR correction. *A. muciniphila* is known to improve MetS in mice^21,22^ and its abundance is negatively correlated with obesity and T2D in humans^23–28^.

Moreover, a recent study showed that *A. muciniphila* supplementation in overweight or obese insulin-resistant humans reduced insulinemia and lowered plasma total cholesterol levels compared to placebo treatment^7^. This validates that our analysis successfully identified microbes protective against the MetS. *Lactococcus lactis* is commonly a part of probiotic mixes, some of which have been shown to improve health^29,30^. Similarly, *L. johnsonii* is used as a probiotic and it is important to test its effects on metabolism.

### *L. johnsonii* does not improve body weight gain on HFD

We set out to test whether microbial factors enriched in obesity-resistant mice would promote healthier host metabolic phenotypes under HFD feeding condition, thereby contributing to the heterogeneity in body weight gain and conferring a fitness advantage to the host. Both *A. muciniphila* and *L. johnsonii* were negatively associated with body weight gain, but since the causal link between *A. municiphila* and metabolic phenotypes has been established previously^7,21^, we focused on *L. johnsonii.* To test whether *L. johnsonii* has causative effects on MetS, we supplemented mice with *L. johnsonii* under a HFD feeding condition.

First, we investigated how well *L. johnsonii* oral supplementation would engraft in the murine intestine. To do this, we gavaged mice with *L. johnsonii* (100ul, 6×10^7 CFU), collected stool samples daily, and quantified *L. johnsonii* abundance in the samples by performing qPCR using primers for *L. johnsonii.* The results showed that *L. johnsonii* abundance returned to baseline within three days **(Fig. 2-5a)**.

**Fig. 2-5.**
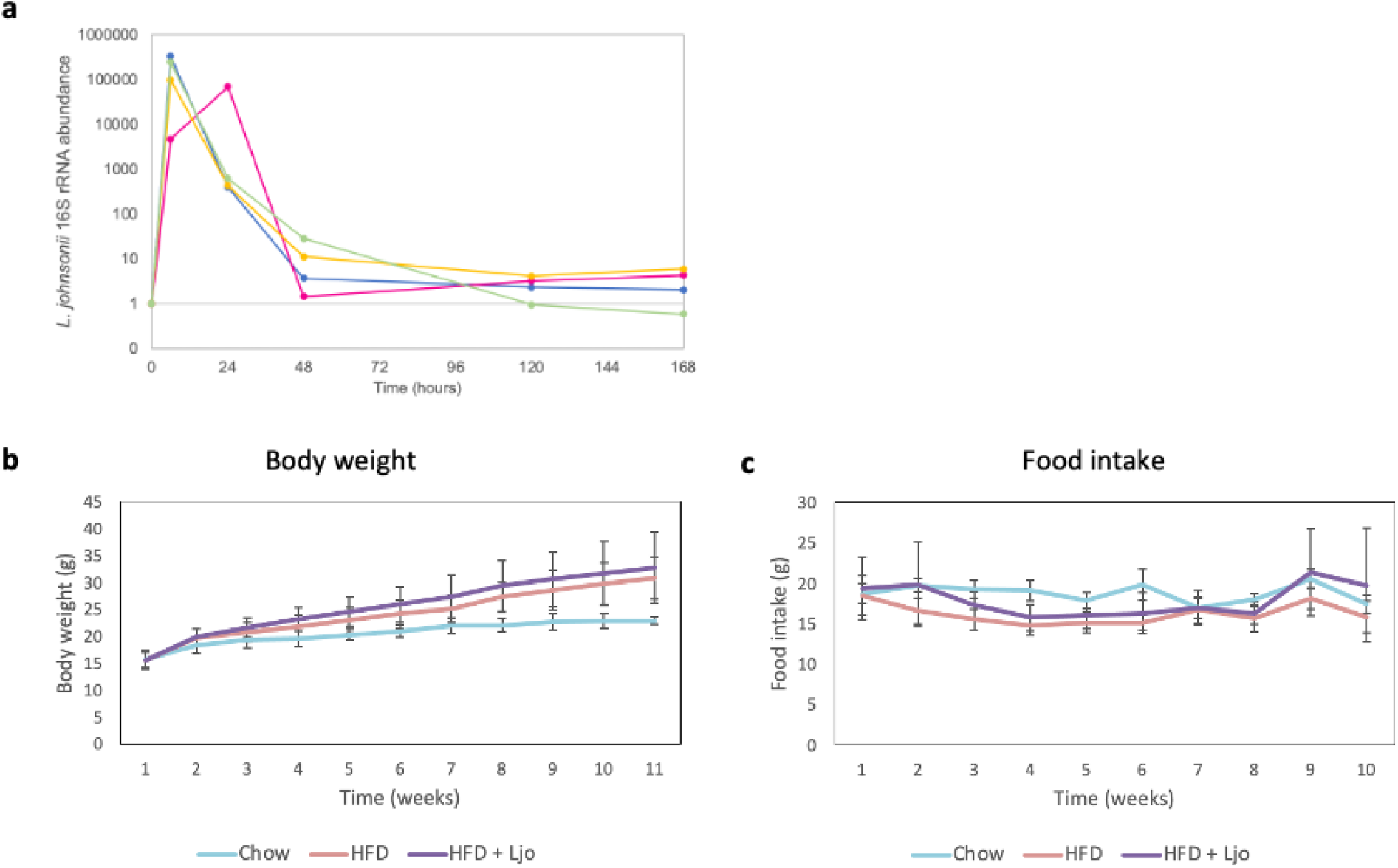
*L. johnsonii* supplementation results. **a,** *L. johnsonii* abundance in the mouse gut over time after supplementation at day 0. Each line represents a mouse. **b-c,** Body weight (**b**) and weekly food intake (**c**) of chow-fed mice, HFD-fed mice, and HFD-fed mice supplemented with *L. johnsonii* (3 times per week) over the course of 10 weeks. Data are shown as average values with error bars representing s.d.

Therefore, we gavaged C57BL/6 mice three times a week with either *L. johnsonii* or PBS control and fed the mice with a HFD for 10 weeks. As control, some mice were kept on a chow diet throughout the experiment. The results showed that there is no significant difference in either body weight gain or food intake between the experimental and the control groups within 10 weeks of HFD feeding **(Fig. 2-5b and Fig. 2-5c)**. Further experiments are needed to identify gut microbial species that play a role in the fitness of its host in the context of MetS and obesity.

### Pathways associated with body weight gain

Moreover, for all mice from all generations, we identified pathways (annotated by HUMAnN2^31^) from stool samples collected after 5 weeks of HFD that are negatively correlated (Pearson correlation) with body weight gain upon 10 weeks of HFD **(Fig. 2-6)**. Notably, almost all of the significantly correlated pathways are from *Akkermansia muciniphila*. These include pathways for coenzyme A biosynthesis, amino acid biosynthesis (lysine, isoleucine, valine), and nucleotide biosynthesis (adenosine, guanosine). Adenosine signaling might play a role in lowering body weight gain as adenosine has been shown to activate brown adipose tissue and recruit beige adipocytes^28^. The activity of brown and beige adipose tissues are associated with lower body mass and fat mass^1,2^, which is consistent with our observation that lower body weight gain is associated with enrichment in adenosine biosynthesis pathways.

**Fig. 2-6.**
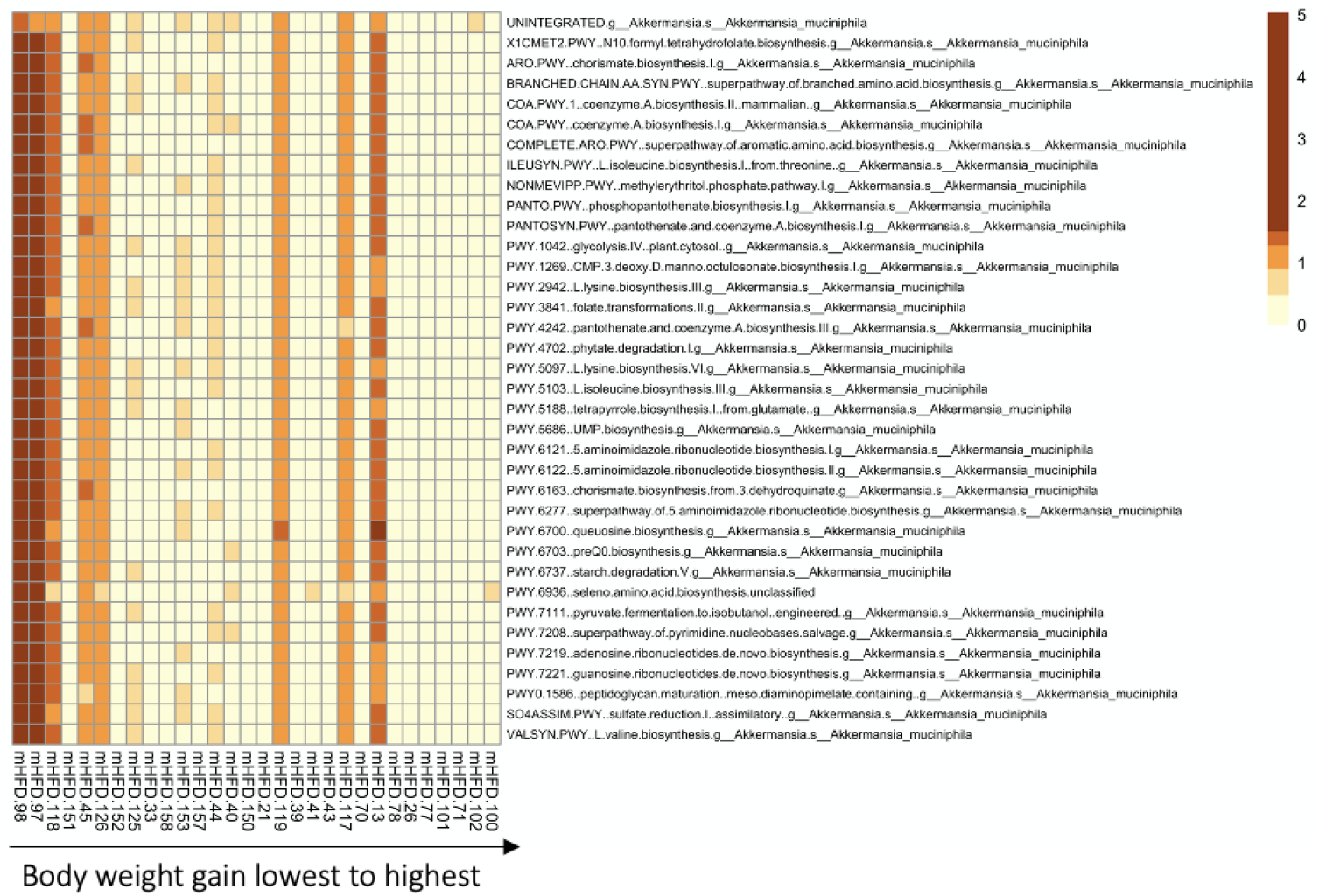
Pathways associated with body weight gain. Microbial pathways were predicted from raw reads using HUMAnN2^31^ from stool samples collected after 5 weeks of HFD treatment. Pearson correlation coefficient between pathway abundance and body weight gain upon 10 weeks of HFD was calculated. Statistically significant pathways were included in the heat map. Each column represents one mouse and each row represents a pathway. Pathway abundances are shown on a log scale and scaled by row.

## Discussion

It is well recognized that obesity negatively affects fertility in both humans^32^ and mice^33,34^. Hence, heritable factors that protect against obesity and MetS and give a reproductive advantage would be present at a higher frequency in subsequent generations. Although human genetics partially explain the heritability of obesity and T2D^35,36^, the human microbiome also plays a significant role. The gut microbiome evolves over a shorter time span, whereas changes in human genetics occur at the population level only after many generations. The gut microbiome is also vertically transmitted across generations^37–40^, therefore, changes in one generation can be passed on to subsequent generations. This raised the question of whether there are inherited gut microbiome factors that play a role in the fitness of the host.

A previous study has shown evidence of the microbiome affecting host natural selection. In this study, exposure to toxic levels of pesticides in *Nasonia* wasps led to changes in the composition of the microbiome that are transmitted to the following generation^41^. After being exposed for multiple generations, the wasps become resistant to the pesticide, and this resistance is mediated by the microbiome. However, it is unknown whether the gut microbiome influences natural selection of its host under obesogenic pressure.

Our unique study design enabled us to test this hypothesis. Since C57BL/6 mice are highly inbred, non-genetic factors are anticipated to be the primary contributors to variation in susceptibility to obesity when fed a high-fat diet. In this study, we show that C57BL/6 mice fed a high-fat diet (60% fat) over four generations develop resistance to obesity and MetS. Unexpectedly, the mice were increasingly leaner and more glucose and insulin sensitive across generations. This phenomenon was due to the most obese mice not yielding progenies, whereas the leanest mice successfully reproduced. Their offspring were also resistant to obesity. This suggests that natural selection was observed. To test whether natural selection is observed in mice under extreme HFD conditions, the study needs to be repeated with a larger N, and breeding challenges should be taken into account when estimating the number of mice needed for the experiments.

Previous studies in which mice were fed with HFD for multiple generations resulted in more severe MetS across generations, which is the opposite of what we observed. This is likely because previous studies used an HFD formulation with lower fat percentage^14,16^ or only fed one of the parents HFD^13,15^. In our study, we applied a stronger selective pressure by feeding both parents 60% fat HFD for 4 generations, therefore only the fittest individuals were able to produce offspring and inherit their microbiome. We believe this strong pressure may better model metabolic phenomena in the current obesity epidemic.

We sought to identify microbial factors enriched in obesity-resistant mice that might promote healthier host metabolic phenotypes under HFD feeding, thereby contributing to the heterogeneity in body weight gain and providing an adaptive advantage to the host. Our analyses revealed that body weight gain was negatively correlated with the abundance of *Lactobacillus johnsonii* and *Akkermansia muciniphila*, and positively correlated with *Lactococcus lactis* abundance. However, mice supplemented with *L. johnsonii* did not exhibit any effects on body weight gain under HFD feeding conditions compared to control mice. Further studies are necessary to validate the role of other microbial factors in the heritability of obesity, MetS, and T2D.

## Materials and Methods

### Mice

Animal research was approved by the Joslin Diabetes Center Institutional Animal Care and Use Committee. All experiments were performed in compliance with all of the ethical regulations.

Male C57BL/6J mice were used throughout the study. Female mice were used for breeding but not used for experiments to avoid stressing mothers and the potential for confounding factors introduced by fluctuation in metabolic phenotypes during pregnancy and nursing. Mice were bred for four generations. They were fed with either a high-fat diet with 60% fat, 20% carbohydrate, and 20% protein (Research Diets D12492) or a chow diet with 10% fat, 70% carbohydrate, and 20% protein (Research Diets D12450J). Experimental mice were fed a HFD for 10 weeks and reversed to chow for 6 weeks, after which the mice were sacrificed. Control mice were kept on a chow diet throughout the experiment for four generations.

Body weight, food intake, and random fed blood glucose levels were measured weekly. At the end of 10 weeks of HFD (pre-reversal) and at the end of 6 weeks of diet reversal to chow (post-reversal), we performed intraperitoneal insulin tolerance test (ip-ITT) to measure insulin resistance and intraperitoneal glucose tolerance test (ip-GTT) to quantify glucose intolerance. At the time of sacrifice, we collected blood, pancreas, brown adipose tissue, and major insulin-sensitive tissues (white adipose tissue, muscle, and liver) to assess metabolic changes in the host. Tissues were frozen in liquid nitrogen directly for qRT-PCR. Parts of the tissues were collected for histology.

### *Lactobacillus johnsonii* isolation and growth

To isolate *L. johnsonii* strains, mouse feces was first suspended in 1x PBS by repeated pipetting, then streaked onto pH 6.8 MRS 1% agar plates (BD). Plates were incubated ~48hrs at 37°C aerobically or anaerobically and colonies were analyzed using MALDI-TOF Biotyper analysis (Bruker). Isolates identified as *L. johnsonii* were then sequenced across the 16S rRNA gene using primers 5’-GCGAGCTTGCCTAGATGATT-3’ and 5’-CAGGTGTTATCCCAGTCTCTTG-3’ to confirm MALDI-TOF identification.

To generate material for gavage, *L. johnsonii* isolates were cultured in pH 6.8 MRS media (BD), incubated overnight at 37°C aerobically without shaking. Cultures were pelleted at 6000g for 10 minutes, then resuspended in an equal volume of 1x PBS. Resuspended pellets were then centrifuged again at 6000g for 10 minutes and resuspended roughly 100x concentrated in PBS and flash frozen. CFU/mL was determined by plating 10μL spots of diluted cultures onto MRS agar plates and counting colonies. Stocks were then diluted to 10^10 CFU/mL prior to gavage.

### *Lactobacillus johnsonii* supplementation

To evaluate the engraftment efficacy of *L. johnsonii* in the mouse gut after supplementation, six male mice from two different litters were each gavaged with 100ul of 5×10^10 CFU/mL bacteria. Mice were single caged throughout the experiment. To monitor *L. johnsonii* abundance over time, stool samples were collected prior to gavage, 6 hours post gavage, and 1, 2, 3, 5, and 7 days post gavage. DNA was extracted from stool samples using a phenol-chloroform protocol as previously described^42^ and *L. johnsonii* abundance was measured with qPCR, which was performed with a SYBR green master mix (ThermoFisher Cat#4309155) and the following primers.

*L. johnsonii* consensus primers:
F: TTTCTGGATCAGCTACAATTTCATCAC
R: CAGAATATGTAAATGTTTCTCACTTCCG

Universal 16S rRNA sequencing primers:

F: GTGYCAGCMGCCGCGGTAA
R: GGACTACNVGGGTWTCTAAT

To determine the effects of *L. johnsonii* supplementation on body weight gain under HFD feeding condition, C57BL/6 mice (6 male mice per group) were gavaged three times a week with either *L. johnsonii* (100uL of 5×10^10 CFU/mL) or PBS and fed with either a HFD or a chow diet for 10 weeks. Body weight and food intake were recorded every week. Mice were individually caged throughout the study.

### Glucose tolerance test

Mice were fasted for 5 hours then administered with glucose solution (2ml per kg of body weight) via intraperitoneal injection. Blood glucose levels were measured using a glucometer (Infinity Blood Glucose Monitoring System) prior to glucose administration and 15, 30, 45, 60, 90, and 120 minutes after injection.

### Insulin tolerance test

Mice were fasted for 2 hours then administered with 1U per kg of body weight insulin (Eli Lilly Humulin R 100U/ml) via intraperitoneal injection. Blood glucose levels were measured using a glucometer (Infinity Blood Glucose Monitoring System) prior to insulin administration and 15, 30, 45, and 60 minutes after injection.

### DNA extraction, library preparation, and shotgun metagenomic sequencing

Faecal samples were collected using sterile forceps and frozen at −80 degrees Celsius until DNA extraction. DNA was extracted from stool samples using either the ZymoBIOMICS DNA Miniprep Kit (D4300) or a phenol-chloroform protocol as previously described^42^. DNA concentration was measured using a Qubit 3.0 with the High Sensitivity DS DNA assay (ThermoFisher). Purity of DNA was determined using a NanoDrop Spectrophotometer. Sequencing libraries were prepared according to a previously published protocol^43^. DNA concentration was again measured using a Qubit 3.0 with the High Sensitivity DS DNA assay (ThermoFisher). Shotgun metagenomic sequencing was performed in 2 × 150-bp paired-end format on the Illumina HiSeq 4000 platform.

### Taxonomic analysis

Microbial taxa were profiled from the raw reads using either MetaPhlAn2 (v.2.7.5)^18,20^ or Kraken 2^19^ and Bracken^20^ with default settings. Alpha diversity was assessed by counting the number of species present in each sample. Pearson correlation coefficient was calculated using the R function cor.test from the stats package (v.3.6.2).

### Pathway analysis

Microbial pathways were predicted from the raw reads using HUMAnN2^31^. Pearson correlation coefficient was calculated using the R function cor.test from the stats package (v.3.6.2).

